# The amyloid concentric β-barrel hypothesis: Models of amyloid beta 42 oligomers and annular protofibrils

**DOI:** 10.1101/2021.08.03.454959

**Authors:** Stewart R. Durell, Rakez Kayed, H. Robert Guy

## Abstract

Amyloid beta (Aβ) peptides, a major contributor to Alzheimers disease, occur in differing lengths, each of which forms a multitude of assembly types. The most toxic soluble oligomers are formed by Aβ42; some of which have antiparallel β-sheets. Previously, our group proposed molecular models of Aβ42 hexamers in which the C-terminus third of the peptide (S3) forms an antiparallel 6-stranded β-barrel that is surrounded by an antiparallel barrel formed by the more polar N-terminus (S1) and middle (S2) portions. These hexamers were proposed to act as seeds from which dodecamers, octadecamers, both smooth and beaded annular protofibrils, and transmembrane channels form. Since then, numerous aspects of our models have been supported by experimental findings. Recently, NMR-based structures have been proposed for Aβ42 tetramers and octamers, and NMR studies have been reported for oligomers composed of ~ 32 monomers. Here we propose a range of concentric β-barrel models and compare their dimensions to image-averaged electron micrographs of both beaded annular protofibrils (bAPFs) and smooth annular protofibrils (sAPFs) of Aβ42. The smaller oligomers have 6, 8, 12, 16, and 18 monomers. These beads string together to form necklace-like bAPFs. These gradually morph into sAPFs in which a S3 β-barrel is shielded on one or both sides by β-barrels formed from S1 and S2 segments.

## Introduction

The quest to determine precise molecular mechanisms by which amyloid beta (Aβ) peptides wreak havoc on the human brain can appear hopeless. These peptides are shapeshifters; they assume countless forms that are often present simultaneously and likely have partially occupied secondary structures (see Deleanu et al.^1^ and Urban et al.^2^ for recent reviews). Numerous factors affect Aβ assemblies; e.g., ions and heavy cations, lipids, concentration, time, method of isolation and preparation, location in the body, initial seed conformation, length of the peptide, and oxidation. Also, they appear to interact with up to twenty receptors^3^, and with several other types of amyloid-forming peptides.

It remains unclear which of their many guises and interactions are the culprits. Much attention has focused on the most visible and stable forms: fibrils within the amyloid plaques that are the hallmark of Alzheimer’s Disease (AD). Their stability has allowed the molecular structure of some forms to be determined experimentally. However, fibrils come in multiple forms: some have U-shaped monomers^4–7^; others have S-shaped monomers^5,8^, some have two-fold symmetry along their long axis and others have three-fold symmetry^5,8^; and most have in-register parallel β-sheets, one has an elongated plus a β-hairpin conformation^9^, and at least one highly toxic mutant forms antiparallel β-sheets^4–7,10^. However, evidence is increasing that much smaller assemblies, called oligomers, are more detrimental (reviewed in^11–13^) and that the longer of the two major forms, Aβ42, is the most toxic^12^. A subset of the oligomer school contends that Aβ oligomers perturb neuronal signaling and eventually kill neurons by forming transmembrane ion channels^13–20^.

Many, perhaps most, large Aβ assemblies reflect their origin: i.e., the final structure depends upon the ‘seed’ structure from which it has grown^21^. Previously we attempted to address these issues by developing atomically explicit models of the structures of Aβ42 hexamers, dodecamers, annular protofibrils, and an ion channel^22,23^. The starting point for our models was the hypotheses that Aβ42 hexamers can adopt a concentric antiparallel β-barrel structure with a hydrophobic core β-barrel, formed by the last third of the sequence (S3), surrounded by a more hydrophilic β-barrel formed by the first third (S1) and middle third (S2) of the sequence. In these models all monomers have well defined identical conformations and interactions with other monomers.

Several aspects of our hexamer model were unprecedented: (1) six-stranded antiparallel beta-barrels had never been reported. However; Laganowsky *et al.*^24^ have since found that a segment from an amyloid-forming protein, alpha B crystalline, indeed has the six-stranded antiparallel β-barrel motif (which they call Cylindrin), and Do *et al.*^25^ found that several eleven-residue peptides with the sequences of portions of S3 that includes methionine also form this motif. Their calculations confirm our findings that the presence of glycine facilitates packing of aliphatic side chains (especially methionine) in the interior of the barrel. The importance of these residues is supported by findings that mutation of Gly33 to Ala^26^ or oxidation of Met35^27^ reduces toxicity and alters oligomerization of Aβ42. (2) Our β structures were antiparallel whereas all known Aβ fibril β-structures were parallel. However, since then an Iowa mutant responsible for some forms of early onset AD has been shown to form fibrils with antiparallel β-sheets^10^. More important, recent experiments indicate that some Aβ42 oligomers do have an antiparallel β secondary structure that is similar to that of OMPA (an antiparallel β-barrel channel)^28–30^, and NMR studies of tetramers, octamers^31^, and 150 kDa oligomers^32^ indicate that S3 β-strands are antiparallel. Also, antiparallel oligomers are more toxic than those with parallel structures^28,30^. (3)Concentric β-barrels had never been observed when we proposed the structures. But recent studies have found that the channel-forming toxins Areolysin^33^ and Lysenin^34^ do, in fact, contain concentric β-barrels. (4) There was no experimental evidence supporting our proposal that the S1 segments form a β-strand or possibly a β-hairpin. However, subsequently two fibril structures with S-shaped monomers that include S1 have been solved (one with 3-fold symmetry^5,8^ and one with two-fold symmetry^4^). In both, the S1 and S2 segments comprise a parallel β-sheet with a bend near the center of S1. We model S1 in two basic conformations: as a continuous β-strand from residues 2-13, and as a β-hairpin with residues A2-H6 forming the first strand (S1a) and residues Y10-H13 forming the second strand (S1b). The β-turn of the hairpin occurs at residues with a high propensity for turn and coil conformations (D7-S8-G9)^35^, but the exact location and direction of the turn varies among the models. Although often ignored by modelers, numerous findings indicate that alterations within S1 segments affect the toxicity of Aβ and that some mutations within S1 are pathogenic (see ^8^ and ^36^ for reviews). Banchelli et al.^37^ found that Cu^++^ causes formation of dimers by binding between His6 and His13 or His14 of two Aβ monomers. They concluded that at least portions of adjacent S1 segments are antiparallel (consistent with some of our models) rather than in-register parallel. Also, Tyr10 side chains can cross-link under oxidizing conditions to form dimers^38^, indicating that they are proximal in some oligomers. (5) There was no experimental evidence that Aβ42 forms β-barrels. However, Serra-Batiste *et al.*^19^ have recently discovered membrane-mimicking conditions under which Aβ42, but not Aβ40 peptides, form a well-defined β-barrel composed of only two monomeric conformations. These results support our proposal that Aβ42 channels contain well-ordered β-barrels resembling those we proposed for oligomers and annular protofibrils.

Experimental constraints used in developing the models presented here were derived primarily from negatively stained electron micrographs of annular protofibrils. Two types of Annular ProtoFibrils have been reported: beaded APFs (bAPFs) that resemble necklaces formed by a string of beads, and smooth APFs (sAPF) that resemble smooth rings^39^. Portions of electron micrographs used in this study are shown in Fig. 1. These APFs form in the presence of hexane, with bAPFs forming initially from oligomers, then gradually transforming into sAPFs. Also, we have incorporated results of recent solid-state NMR studies of oligomers^31,32^ in our latest models.

**Figure 1.**
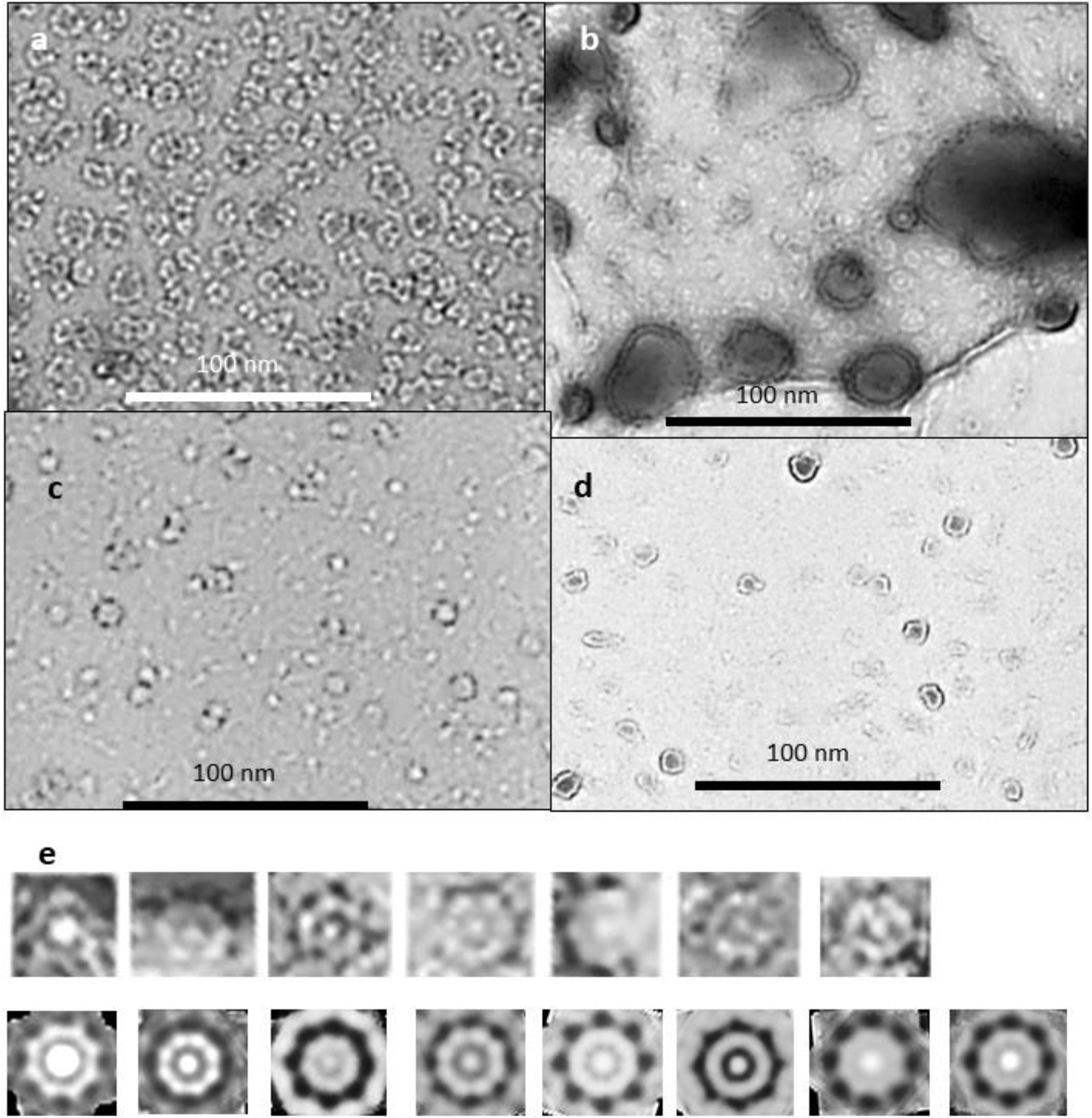
Negative stained EM images of beaded (a) and smooth (b, c, & d) APFs (originals provided by Rakez Kayed). See^39^ for methods. (e) Example of how averaged images were developed. These images are from EM of (b) for one size of WCsAPF that has 8-fold radial symmetry. Images of individual APFs were copied into Photoshop and intensity and contrast were adjusted to make their characteristics more visible (top row). Each of these was averaged with radial symmetry (bottom row). All radially averaged images of similar size and symmetry were aligned and averaged to obtain the final image for each specific sAPF (last image of the bottom row).

## Methods

Diameters of β-barrels comprising these models were calculated from the number of strands (N) and sheer number (S) (related to strand tilt) of the barrels (see Murzin *et al*.^40^ for early theory and Hayward and Milner-White^41^ for an extension and modification of this theory to radially symmetric β-barrels and β-helices). The latter analysis does not include the P2 2-fold perpendicular symmetry present in our models. These structures have axes of 2-fold symmetry perpendicular to the radial axis of symmetry and intersecting it at the assembly’s center of mass. This feature constrains symmetric models with concentric β-barrels because all barrels must have the same axes of symmetry. The gap distance between the walls of the adjacent barrels were constrained to be between 0.6 and 1.2 nm^42^, depending primarily on the side chain sizes. We favor models in which interacting pleats of adjacent β-barrels fit between each other in a manner that reduces clashes among side chains, and in which all pleats that intersect the axes of 2-fold symmetry have the same inward or outward orientation in all concentric β-barrels. We call this type of interaction “Intermeshing Pleats^42^.” We selected models that maximize hydrogen bonds, salt bridges, interactions among aromatic side chains, burial and tight packing of hydrophobic side chains that are highly conserved among Aβ homologs, and aqueous solvent exposure of hydrophilic side chains, especially for positions that are hypervariable among homologs. Residues with polar side chains and/or that have a high propensity for turn or coil secondary structure^35^ and that contain proline in some homologs were favored for connecting loops. The final constraints were experimental: the sizes, shapes, molecular weights, and secondary structures of assemblies as determined by EM images of annular protofibrils and other studies, and distance constraints based on NMR studies of tetramers, octamers, and 32mers.

Photoshop^43^ was used to analyze APF images. Isolated beaded APFs (Fig. 1a) were copied and grouped according to the number and size of beads. Radial image averaging was applied to relatively symmetric bAPFs in which all beads had the same approximate size. The radially averaged images of the same size and shape were superimposed and averaged to obtain the final image. A similar process was used to analyze sAPFs but with the additional constraint of placing the sAPFs into three categories: those with white centers and periodic dots on the perimeters (WCsAPF), those with dark centers and periodic dots (DCsAPF), and those with super smooth outer rings without periodic dots and dark centers (SsAPF). The first two categories of images were then averaged radially with the symmetry based primarily on their diameters and the number and spacing of the perimeter dots. Averaged images that were similar in size and characteristics were then aligned to obtain images averaged over multiple sAPFs (Fig. 1e). When the symmetry was ambiguous numerous symmetries were tried. If a specific symmetry and diameter resembled those obtained from less ambiguous sAPFs the image was included in the multiple sAPF averaging process; otherwise, it was discarded.

Atomic-scale structures were generated with an in-house program and illustrated using the Chimera program^44,45^.

Sequences of 2500 Aβ homologs were collected and aligned using the Blast program^46^.

## Results and Discussion

### Oligomers and beaded APFs

The first method of evaluating the sizes of the beads was to copy individual images of isolated beads in both bAPF and sAPF micrographs, classify them according to their sizes, superimpose the images in each category, and then use image averaging to obtain a single image for each size (first two rows of Fig. 2). We identified five sizes of beads. The major difficulty with this method is that the outer perimeter is not clear cut. Although isolated images of the smallest category were difficult to identify in the bAPF micrograph, they were prevalent in some sAPF micrographs.

**Figure 2.**
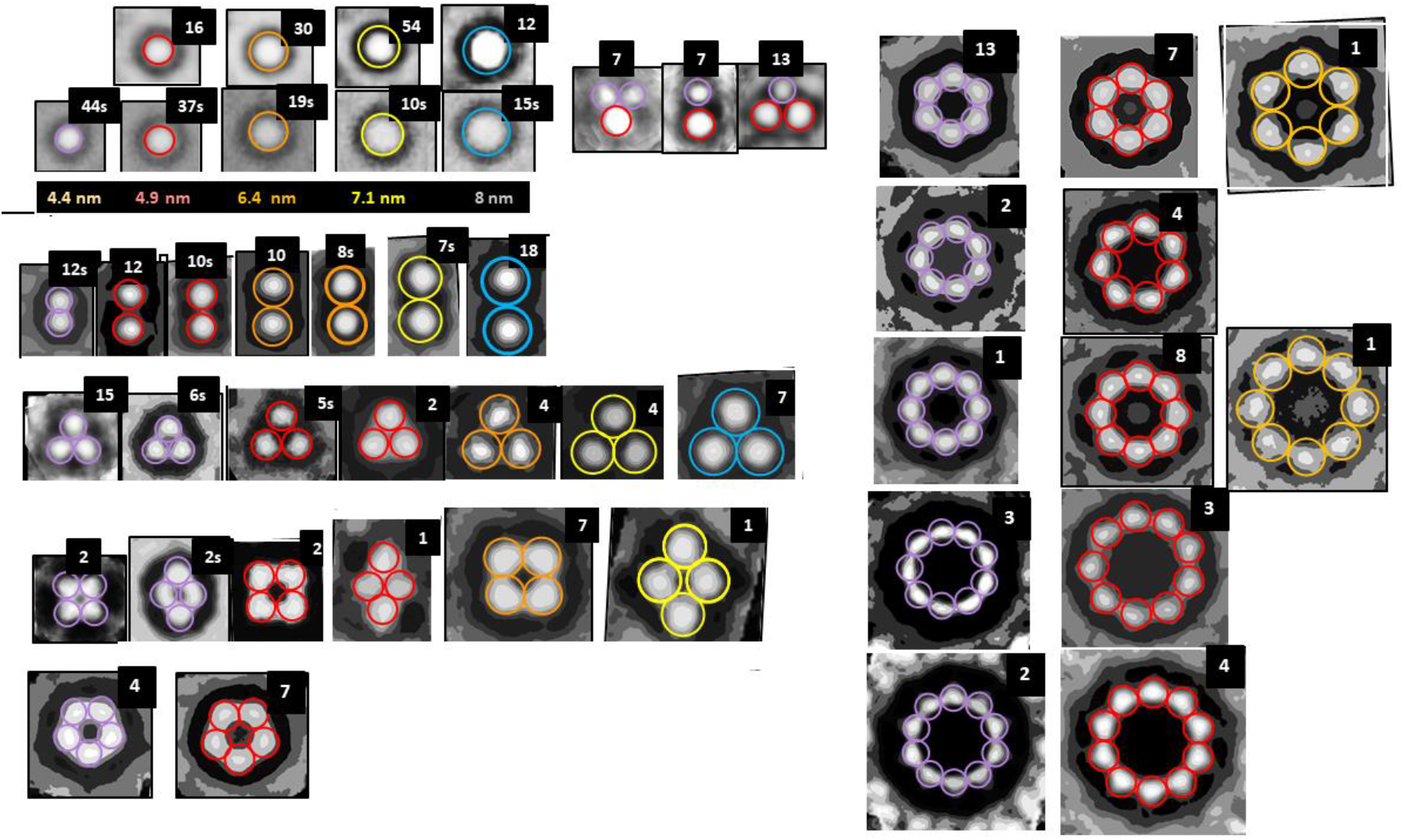
Averaged images of isolated and symmetric assemblies of beads (oligomers). The colored circles have the diameters indicated below the second row. The number of images used to develop the averaged images is in the upper right corner; when followed by a “S” the images were made from one of the sAPF micrographs. There appears to be five sizes of beads with diameters ranging from about 4 to 8 nm. The first two rows on the left-half show isolated beads from both types of micrographs followed by three examples of small clusters composed of the two smallest-size beads. The remaining rows shows symmetric assemblies composed of 2 to 10 beads. Beads within each illustrated bAPF all have the same size, but the bead size varies among bAPFs that have the same number of beads. The two smallest sizes occur most often for bAPFs with five or more beads. The “cutout” facility of Photoshop was applied to most images to help identify centers of the beads.

The second method was to approximate distances between beads of bAPFs. We first identified relatively symmetric bAPF composed of the same size of beads. We used rotational image averaging to obtain symmetric averaged images, and then superimposed images of the same size and shape to obtain a single averaged image. We excluded bAPFs from the analysis if they were relatively asymmetric, had substantial gaps between beads, were composed of different sized beads, contained beads that appear to have merged, and/or had an ambiguous number of beads. The advantage of this approach is that the diameter of the beads can be approximated from the distance between the centers of adjacent beads, which can be measured more precisely than the location of the outer edges of isolated beads. The remaining rows in Fig. 2 are images of assemblies of 2-10 beads. Sizes of the beads are color coded as in the first two rows.

### Models of oligomers that correspond to “beads”

The next challenge was to generate structural models consistent with these bead sizes. The following features are common to all small non-APF oligomer (bead) models presented below.

a. The number of monomers, M, of an oligomer is an integer multiple of six or eight. Models presented here are only for assemblies of six or more monomers. Hexamers, dodecamers, and octadecamers are based on results of cross-linking studies^47^. Octamers are based on solid-state NMR studies^31^.
b. All models have two or three concentric β-barrels. Six or eight antiparallel S3 β-strands comprise the innermost β-barrel.
c. All have 2-fold perpendicular symmetry and 3-fold or 4-fold radial symmetry.
d. All adjacent β strands are antiparallel.
e. Except for the octadecamer, all putative β-strands are components of β-barrels. S1 can be either a S1a-S1b β-hairpin or a single β-strand.
f. S1a-S1b β-hairpins and S1 β-strands are on the surface; hydrophilic sides of the strands are exposed to water while hydrophobic side chains on the opposite side interact with S3 or S2.
g. The vast majority of side chains buried in the interior of the assembly are hydrophobic, the few buried polar side chain atoms from the outer strands can form H-bonds with other groups.
h. Electrostatic interactions are also favorable; almost all charged groups form salt-bridges and almost all polar backbone atoms form H-bonds.

Concentric β-barrel models of oligomers presented in the text are not unique; some alternatives with differing details are described in the supplement. Resolutions of EM images and molecular modeling methods are not sufficient to identify the best of these alternatives. However, all are consistent with the hypothesis of concentric β-barrel assemblies.

The smallest β-barrel oligomer modeled here is a hexamer (Fig. 3). S3 strands form a hydrophobic core as a six-stranded antiparallel β-barrel. Although six stranded β-barrels are rare, this one is made possible by the absence of side chains at the inward-pleated G33 and G37 residue backbones, and the flexibility of the M35 side chains that fit next to these glycines. The S3 core of the hexamer is surrounded by an 18-stranded antiparallel β-barrel formed by S1a, S1b, and S2 strands. The illustrated hexamer has an outer diameter of ~4.4 nm (Fig. 3a) assuming an outer diameter 1.0 nm greater than the diameter of the outer β-barrel’s wall. (An alternative model with a S/N ratio of 2/3 has a diameter of ~4.1 nm (see supplement Fig. S1)). Side chains at the axes of 2-fold symmetry intermesh; i.e., V18 of S2 and V36 of S3 (green circle) are both oriented outwardly, and F4 of S1a and M35 of S3 (purple circle) are both oriented inwardly.

**Figure 3.**
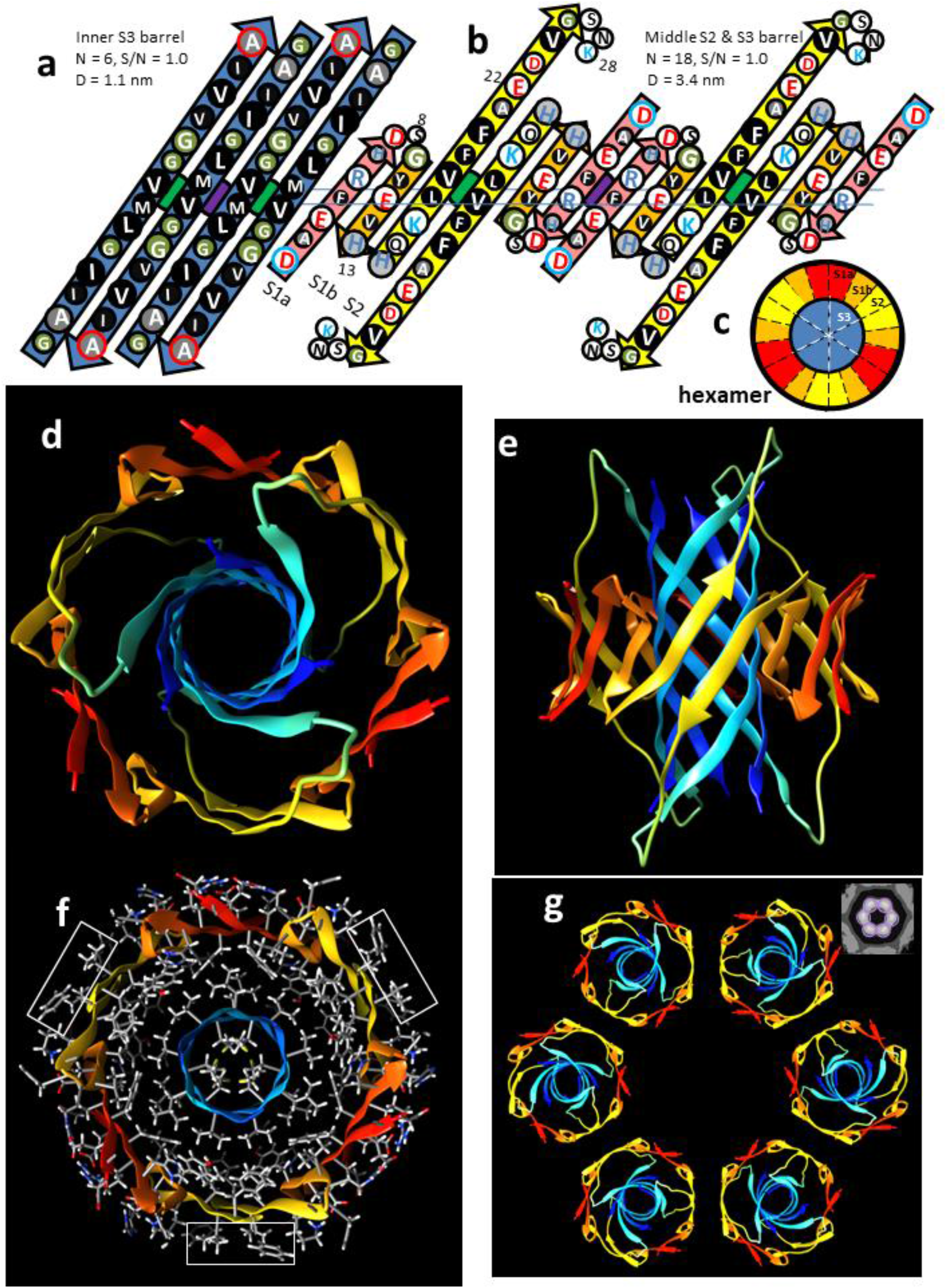
Hexamer concentric β-barrel model. (a & b) Flattened representation of four subunits viewed from the exterior as if the barrel were split open on the back side and spread flat. The axes of 2-fold perpendicular symmetry are indicated by green and purple circles behind the β-sheets, and the plane containing these axes by the horizontal line. The arrows represent β-strands colored by segment. The circles represent side chains; those oriented outwardly are larger than those oriented inwardly. They are colored by residue type: white = positively charged (blue letter), negatively charged (red letter), or uncharged polar (black letter), green = ambivalent Gly, light gray = ambivalent His (blue letter) or Thr (black letter), dark gray - white letter = slightly hydrophobic Ala, black - white letter = hydrophobic. The N- and C-termini residues have blue and red outlines to indicate their positive and negative charges. (a) Outer β-barrel; S1a (pink), S1b (orange), and S2 (yellow); parameters are N = 18, S/N = 1.0, D = 3.4 nm. (b) Inner S3 β-barrel; N = 6, S/N = 1.0, D = 1.1 nm. (c) Wedge representation cross-section on the plane containing axes of 2-fold symmetry illustrating relative positions of the β-strands. (d-g) Atomic-scale model. The backbone is illustrated as a rainbow-colored ribbon from red (N-termini) to blue (C-termini). (d) View down the radial axis of the β-barrels. (e) Side view along the 2-fold axis between the yellow S2 strands. (f) Radial cross-section through the central portion with side chains colored by element; hydrogens and carbons are white and gray, polar oxygen and nitrogen atoms are red and blue. Rectangles enclose exposed V18 and F20 side chains. (g) Ribbon representations of six hexamers forming a beaded APF; the insert in the upper right corner is an averaged EM image from Fig. 2.

The exterior surface of the hexamer model has three hydrophobic patches formed by V18 and F20 (rectangles of Fig. 3f) flanked by positively charged K18 and negatively charged E22. These are the most probable interaction regions between APFs in an aqueous solution due to a combination of hydrophobic and electrostatic interactions. Such interactions should be geometrically optimal for hexamers in a bAPF formed from six small beads due to the 2-fold symmetry of the interaction sites (Fig. 3g). In the beaded APF micrograph analyzed here, six-membered APFs dominated the smallest bead APFs; 13 six-membered rings were observed whereas the average number of other small bead bAPFs was only about two (Fig. 2).

Experimental studies indicate that Aβ42 can form dodecamers^48,49^ and octadecamers^50,51^, and that these sizes of oligomers can be isolated from brains of Alzheimer’s patients. Some of these may form when additional monomers bind to hexamers. If so, the S3 strands of the additional Aβ monomers will likely bind in a hydrophobic environment. Site-Directed spin-labeling studies indicate that mobility decreases and rigidity increases within Aβ oligomers from the N-terminus to the C-terminus, and that S3 segments are tightly packed^52^. S1s comprise the most polar, most dynamic, and least conserved of the segments; so much so that in most experimentally-determined structures they are classified as disordered. We hypothesize that S1 segments can be displaced as S2 and/or S3 segments of six additional monomers bind next to S3s and between S2s of the original hexamer (Figs. 4 & 5). The displaced S1a-S1b β-hairpins of the original hexamer may combine with S1a-S1b β-hairpins of the additional monomers to form an outer 24-stranded antiparallel β-barrel with approximate 6-fold radial symmetry (Fig 4), but the outermost barrel or layer is likely to be highly dynamic and difficult to predict precisely.

**Figure 4.**
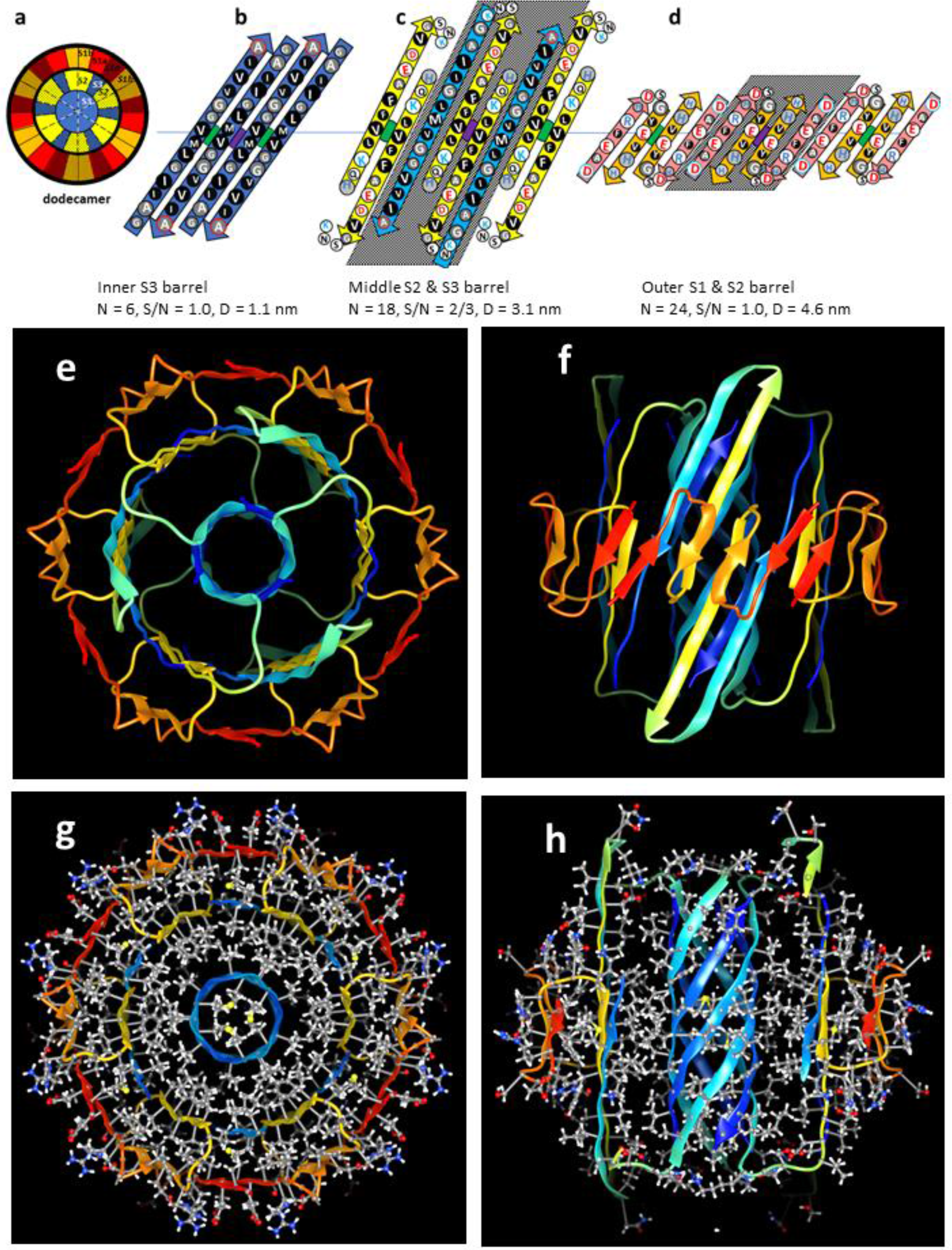
Dodecamer models. Parameters of the barrel are listed below the octadecamer schematics. The color code for the dodecamer strands and residues is the same as in the previous figures. The structures of the core S2 and S3 strands of the hexamer are maintained as additional subunits (stippled), and the core S1a-S1b hairpins (sides of outer barrel) are displaced outwardly to the third barrel. (a) Wedge representations of the central cross-section; peripheral subunits are stippled. (b-d) Flattened schematics of four subunits of (b) the inner 6-stranded S3 β-barrel (same as hexamer), (c) the middle 18-stranded S2 and S3 β-barrel, and (d) the outer 24-stranded S1a-S1b β-barrel. (e-h) Atomic-scale models of the dodecamer. (e & f) Rainbow colored ribbon illustration of the backbone as viewed from the top and side. (g & h) Cross-sections with side chains colored by element viewed along the barrels’ axis and from the side. Note that most buried side chains are tightly packed and hydrophobic (gray carbons, white hydrogens, and yellow sulfurs), and that most surface side chains have polar red oxygen or blue nitrogen atoms with white hydrogens.

**Figure 5.**
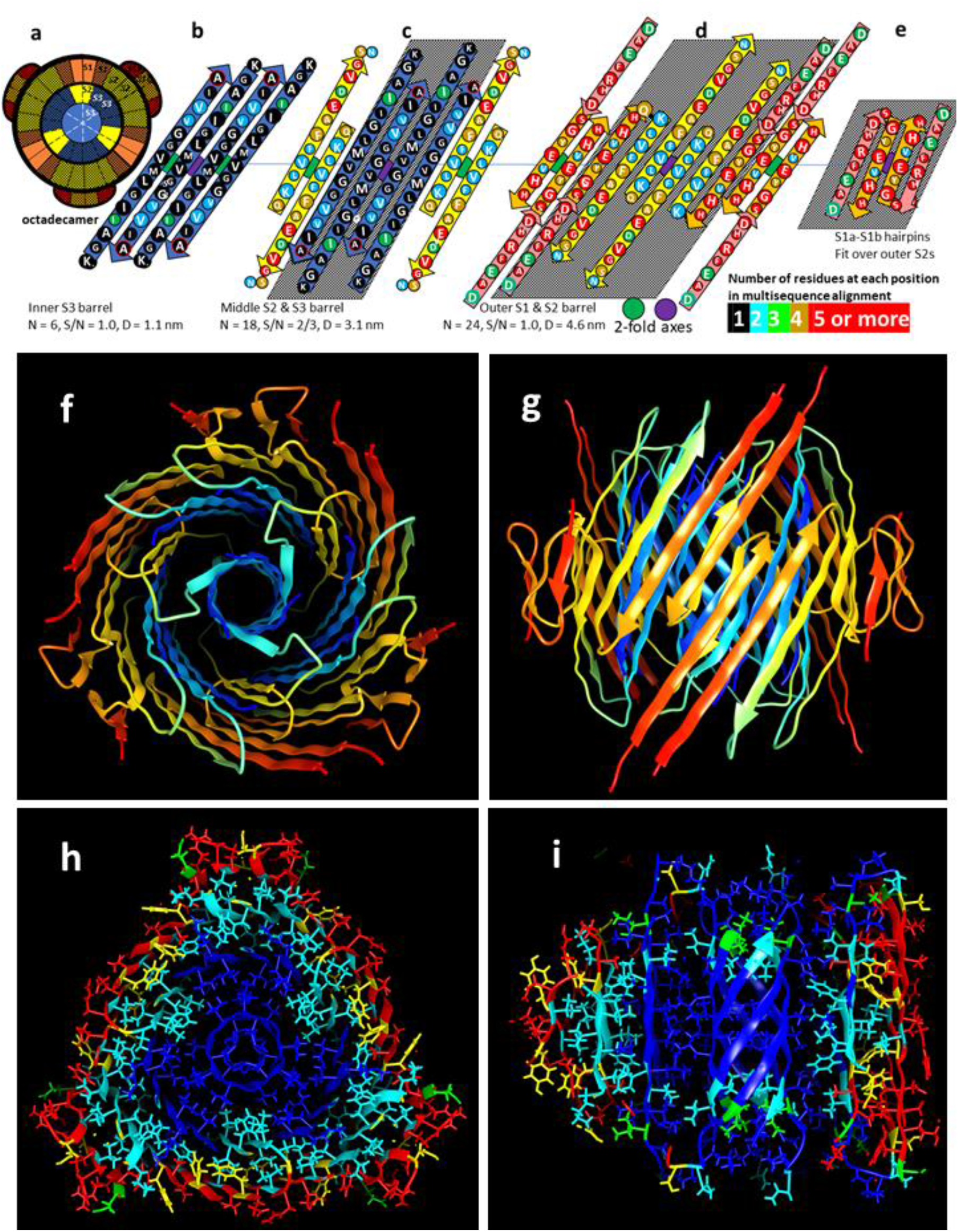
Octadecamer model. (a) Wedge representation of strand positions in the central cross-section; the twelve peripheral subunits are stippled. (b-e) Flattened representation of (b) the core 6-stranded S3 β-barrel, (c) the middle 18-stranded S2-S3 β-barrel; the poorly conserved S1a interacts with the poorly conserved portion near the end of S2 and its structure is likely more disordered than depicted, (d) the outer 24-stranded S1-S2 β-barrel, and (e) two peripheral S1a-S1b β-hairpins. Residues are colored according to the number of residue types at each position of a multi-sequence alignment. (f-i) Atomic-scale models. (f) Top view and (g) side view cross-section of backbone ribbon representation colored by rainbow. (h & i) Model with conserved side chains colored by degree of conservation as in b-e; (h) middle cross-section, (i) side view cross-section.

Aβ homologs comprise a large superfamily present in most vertebrates. Multiple studies indicate that they have a functional role^38,53,54^ including antimicrobial activity^55^. We aligned 2500 homologous sequences from mammals through bony fish (Supplement Fig. S2). Residues 28 – 42 of the S3 segment are incredibly conserved; the sequences are identical in all but three positions and even those substitutions are limited to large alkyl side chains: i.e., I32-V or L, V39-I, and V40-I (Fig. S2). Only one substitution occurs at V12 in S1b, and at K16, L17, V18, F19, and N27 in S2. Only three or four residue types occur at D1 and E3 in S1a, at Y10 in S1b, and at Q15, F20, A21, D23, and S26 of S2. All hypervariable positions where five or more residue types and deletions occur are in S1 or toward the end of S2. The high degree of conservation in S2 and S3 suggest that these proteins adopt a functional conformation or conformations in which at least some S3 segments and conserved portions of S2 are buried within the protein complex, and in which most of S1a is peripheral and likely disordered. Mutagenesis studies indicate that S3 residues I31, I32, L34, V39, V40 and I42 are key to Aβ oligomerization^56^. The octadecamer model of Fig. 5 was developed in part based on the hypothesis that all highly conserved hydrophobic residues form a core structure that is buried in this model. S1a is peripheral as are outwardly oriented side chains of S1b. Side chain packing between barrels is quite tight with few large cavities; all conserved hydrophobic side chains (blue and cyan) and most semi-conserved side chains (yellow and green) are buried, the hypervariable side chain positions (red) are peripheral, backbone H-bonding is extensive, and almost all charged groups form salt bridges.

We suspect that the second largest oligomer (bead) is an octamer. Identification and development of a best model was complicated. An NMR-based model of an Aβ42 structure has been proposed for both tetramers and octamers^31^. For those, each tetramer forms a six-stranded antiparallel β-sheet with 2-fold perpendicular symmetry and two different subunit conformations. Two central S3 strands resemble those of our model; they are antiparallel with a perpendicular 2-fold axis at V36. The remaining portion of these subunits was classified as disordered. S3’s of these subunits are flanked by S3 strands that are oriented in the opposite direction relative to the plane of the β-sheet; *e.g.* side chains of odd numbered residues of the flanking S3 strands are on the opposite side as those of the central S3 strands. S2 strands of these subunits comprise edge strands as part of a β-hairpin with S3. The structure of the S1 region was undetermined. The octamer structure was proposed to be a β-sandwich in which two tetramers related by 2-fold symmetry pack back-to-back. These structures were developed in the presence of detergents, and thus may differ from the APFs analyzed here that were developed in the presence of hexane.

We used atomic-scale modeling to explore the feasibility of octamer structures with concentric β-barrels. After developing numerous models, we concluded that the structure illustrated in Fig. 6 is the soundest that is also consistent with the size of the next to smallest beads, those circled in red in Fig. 2 (See Fig. S3 of the supplement for alternative models). Some aspects of this model resemble those of the hexamer model; all monomers have the same conformation due to radial and 2-fold perpendicular symmetry, and an antiparallel S3 β-barrel with S/N = 1.0 that forms the hydrophobic core. There are however substantial differences in addition to the number of subunits: (1) S3 strands of the 8-stranded core barrel are oriented radially in the opposite direction; i.e., G33, M35, and G35 are on the exterior instead of on the interior. (2) The β-turn linking S1b to S2 centers at H14-Q15 instead of H13-H14 and turns in the opposite direction. (3) S1 is a single continuous β-strand instead of a S1a-S1b β-hairpin, and thus each subunit contributes only two instead of three strands to the outer barrel. (4) The S1 and S2 strands of the outer β-barrel are more tilted; i.e., S/N = 1.5 instead of 1.0. These changes result in the outer barrel having 16 strands and a diameter of 3.6 nm. The gap distance between the two barrels is about the same as for the hexamer model. The pleats intermesh at the perpendicular 2-fold axes: i.e., the outwardly-oriented M35 side chains fit into the pleats behind the outwardly-oriented D7 residues, and the inwardly-oriented F19 side chains fit in the pleats of inwardly-oriented V36 residues. Aromatic side chains of inwardly-oriented F4, Y10, and F19 are adjacent on the same pleat and interact (interactions between aromatic groups tend to be energetically favorable^57^). Locations of G33 and G37 on the exterior of the S3 barrel allow F4, Y10, and F19 side chains to pack next to S3 without major side chain clashes; note that the only outwardly-oriented S3 side chains in the middle cross-section are those of M35 (Fig. 6f). The outer barrel has several inwardly-oriented side chains with polar atoms: i.e., the hydroxyl groups of S8 and Y10, the imidazole ring of H6, and the carboxyl group of D23. These are near enough to one another to form H-bonds and salt-bridges. All positively and negatively charged side chains on the exterior of the outer barrel form salt-bridges. β-barrels in proteins often have eight strands^58,59^. The interior of the 8-stranded S3 barrel is large enough to accommodate the alkyl side chains of I32, L34, V36, and V40 and still leave a narrow pore lined with alkyl groups through its center. Portions of this pore may be occupied by hexane.

**Figure 6.**
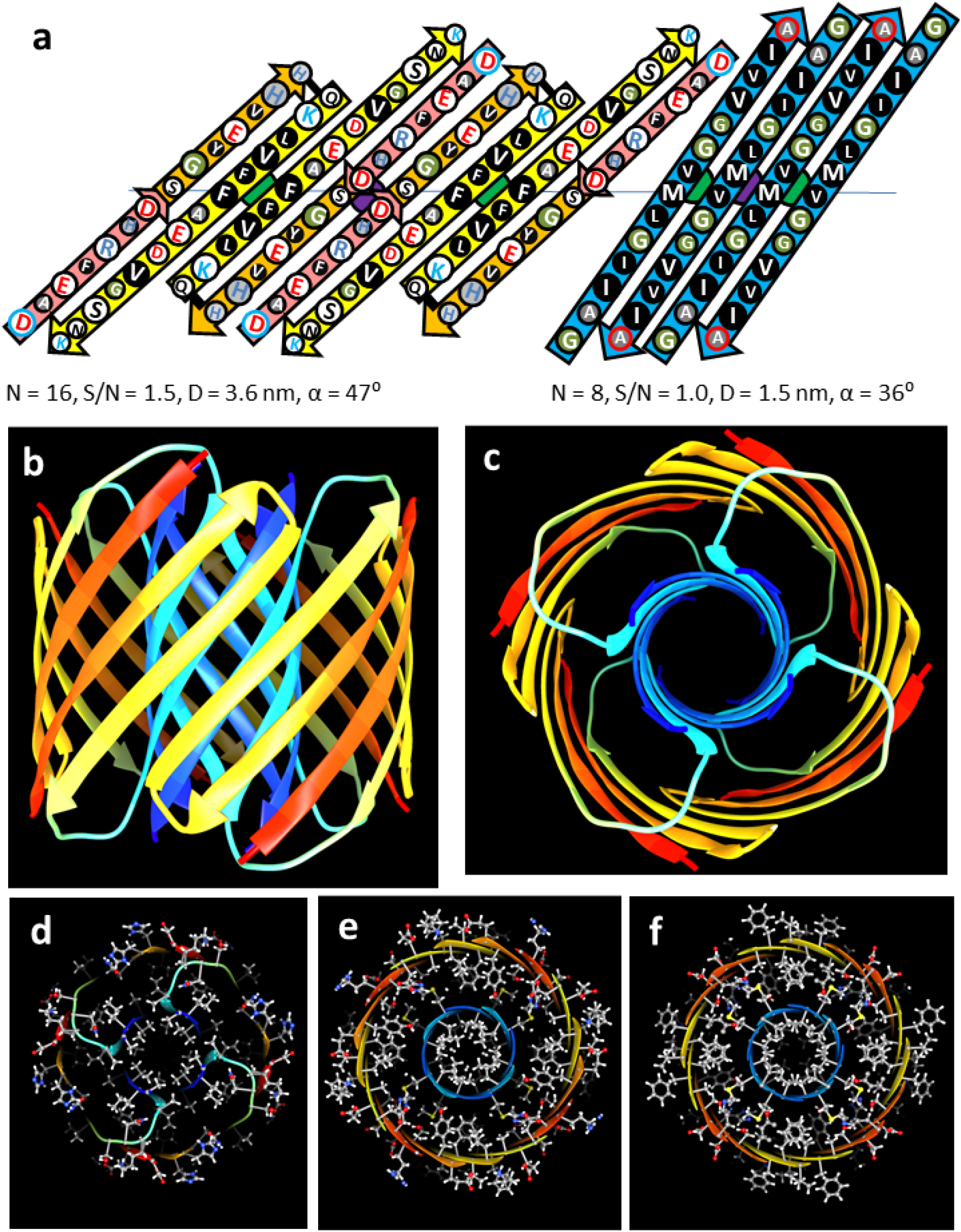
Model of an Aβ42 octamer. Features and colors are like those of Fig. 3. (a) Flattened schematic of four subunits of the outer and inner β-barrels. (b & c) Ribbon representation of the octamer backbone as viewed from the side through a perpendicular 2-fold axis between yellow S2 strands and from the top. (d-f) Cross-sections with side chains colored by element of the top region, the next region down, and the middle region.

### Smooth Annular Protofibrils

Beaded APFs gradually morph into smooth APFs. Most sAPFs in the upper portion of Fig. 1b exhibit concentric rings: an outer dark ring, followed by a light ring, followed by a second dark ring which surrounds a circular light center. These sAPFs resemble cog gears or cog wheels due to evenly spaced dark dots on the perimeter of the outer ring; thus, we call these dots cogs. White-centered cog APFs (WCsAPFs) are densely packed in the top region of the micrograph of Fig. 1b, which appears to be enclosed or covered by a film. Many dark centered sAPFs (DCsAPFs) in other portions of Fig. 1b and in Fig. 1c also have cogs on their perimeters, but they have only an outer dark ring and an adjacent light ring. The micrograph of Fig. 1d is dominated by super smooth APFs (SsAPFs) that have few or no cogs and relatively dark centers.

The concentric rings of WCsAPFs EM images provide the strongest pictorial evidence that Aβ42 peptides can assemble into concentric β-barrel structures (Fig. 7). Radial image averaging should be used cautiously because it can produce the appearance of radial symmetry even when none exists. Selected unaveraged images were included to illustrate that some WCsAPFs, especially the larger ones, exhibit radial symmetry without radial averaging.

**Figure 7.**
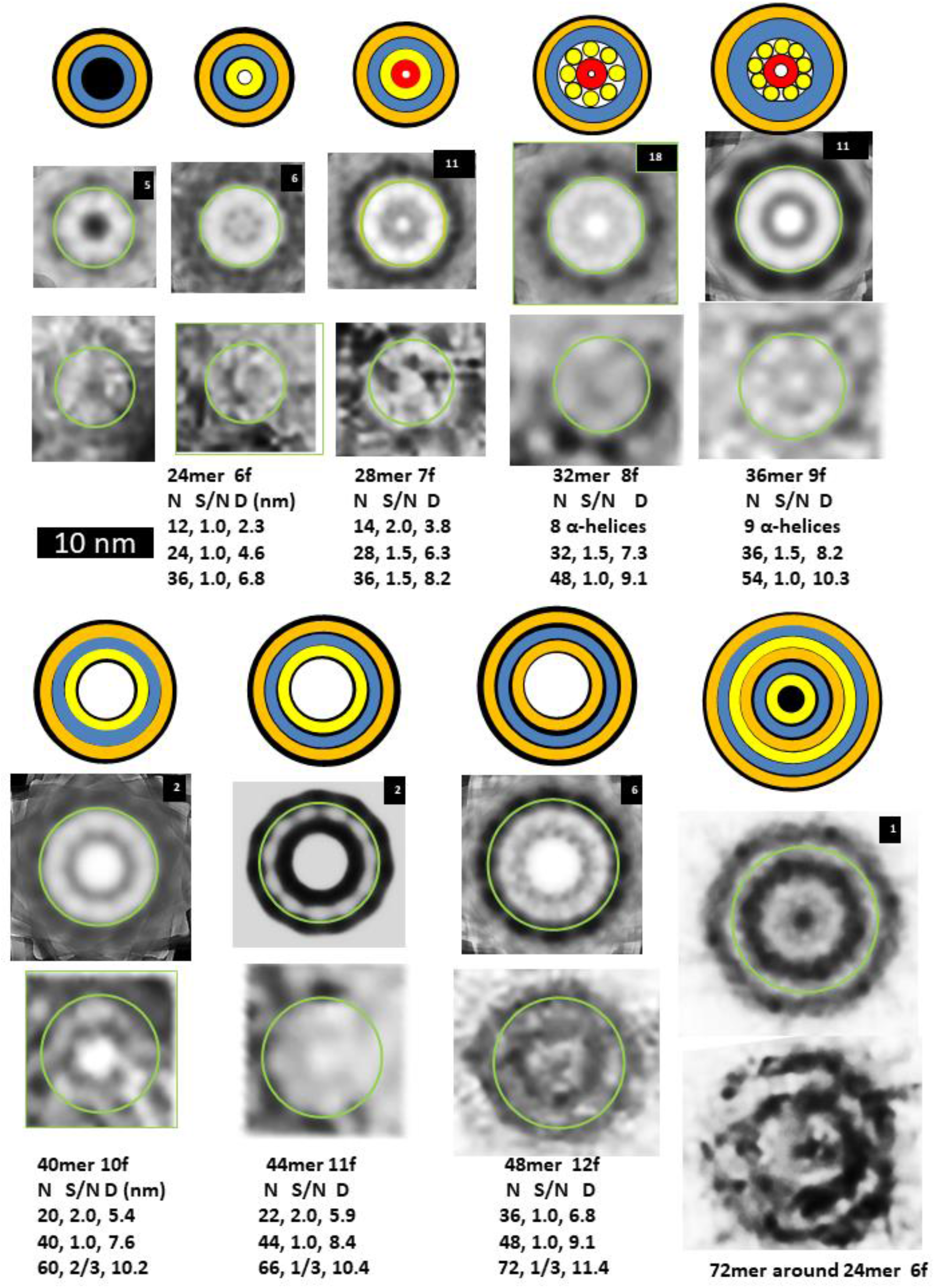
Images of WCsAPFs and schematics of Aβ42 models with three concentric β-barrels. The outer orange rings of the schematics represent the outer β-barrel formed by S1 and S2, the blue ring represents a S3 β-barrel, the inner orange ring represents an inner β-barrel formed by S1 and S2, the inner yellow ring represents a β-barrel formed only by S2 strands, and the tiny red ring represents a highly tentative β-barrel formed by S1a segments. The yellow cylinders of the 32mer and 36mer models represent α-helices formed by S1b-S2 segments. Parameters of the models are listed below the images. The upper rows of images were averaged radially for the number of WCsAPF’s indicated in the upper right corner. The lower row of images are individual sAPFs. The green circles on the images indicate the proposed size and location of boundary between the outer and middle β-barrels. A small DCsAPF (the first image) was included because it also occurs frequently within the same region of the micrograph. The final schematic/EM image is larger and has more concentric rings than the others. The schematic illustrates that the assembly may have six concentric β-barrels with a three-barrel 72mer sAPF surrounding a three-barrel 24mer sAPF (also see Supplement Fig. S5).

WCsAPFs can be classified into seven sizes. Each image was first averaged radially and then all images of the same size and radial symmetry were aligned and averaged (Fig. 1e). The number of images used for each size is indicated in the upper right corner of Fig. 7. We hypothesize that the outer white ring corresponds to a S3 β-barrel, that the number of strands in this barrel is the same as the number of monomers in the assembly, and that the S/N value is either 1.0 or 1.5. These assumptions permit the number of monomers in the assembly to be calculated from the diameter of the white ring (see parameters below the images). Our calculations suggest that the number of monomers increase by four for each size increase. Also, the number of cogs in the outer dark ring increases by one for each size increase. Assuming one radial unit-cell per cog, each radial unit-cell contains four monomers. Thus, these sAPFs may develop and grow from mergers of tetramers. If the tetramer structure of Aβ42 proposed by Cuidad et al.^31^ for S3 strands is approximated in the S3 barrel of the white-centered sAPFs, then the radial unit-cell would be composed of four monomers and contain two subunit conformations. Although an α-helical motif for S1b-S2 of C_in_ may be viable for 36mers and possibly 28mers, size restrictions make it less likely for other white-centered sAPFs. We suspect that C_in_ subunits of the other WCsAPFs form antiparallel inner β-barrels with the strand tilts and involvement of S1 segment changing as the size of the assembly changes, in a manner that keeps the gap distance between the barrels near 1.0 nm.

The final image and schematic of Fig. 7 illustrates the most extreme example of multiple concentric β-barrels. The unaveraged image has a relatively hexagonal shape. Averaging with 6-fold symmetry reveals multiple concentric rings. Fig. S5 of the supplement compares images of putative 24mers and a 72mer to this image, and illustrates details of a model in which a 72mer sAPF surrounds a 24mer sAPF. The central rings of the image resemble the first of the WCsAPF images, the putative 24mers. The outer diameter of the image is consistent with a 72mer sAPF with S/N = 0.5 for the S3 barrel. The model has six concentric β-barrels, but this type of assembly is rare; this is the only image that we observed with these properties.

Like the tetramers proposed by Cuidad et al.^31^, our WCsAPFs models have two monomeric conformations and 2-fold perpendicular symmetry. An antiparallel S3 β-barrel is sandwiched between two cylinders formed by S1-S2 segments. In our models S1a-S1b-S2 β-strands of half the monomers form an outer β-barrel with a structural motif resembling that proposed above for hexamers (Fig. 8). S2 of these monomers connects to the S3 strands for which centrally located V36 side chains are oriented outwardly. We call the conformation of these monomers C_out_. The outer two barrels of C_out_ subunits intermesh at the axis of perpendicular 2-fold symmetry; i.e., the outwardly oriented V18 residues of S2 fit over the outwardly-oriented V36 residues of S3, and inwardly-oriented F4 residues of S1a fit over inwardly-oriented V36 residues of the other S3 conformation, C_in_.

**Figure 8.**
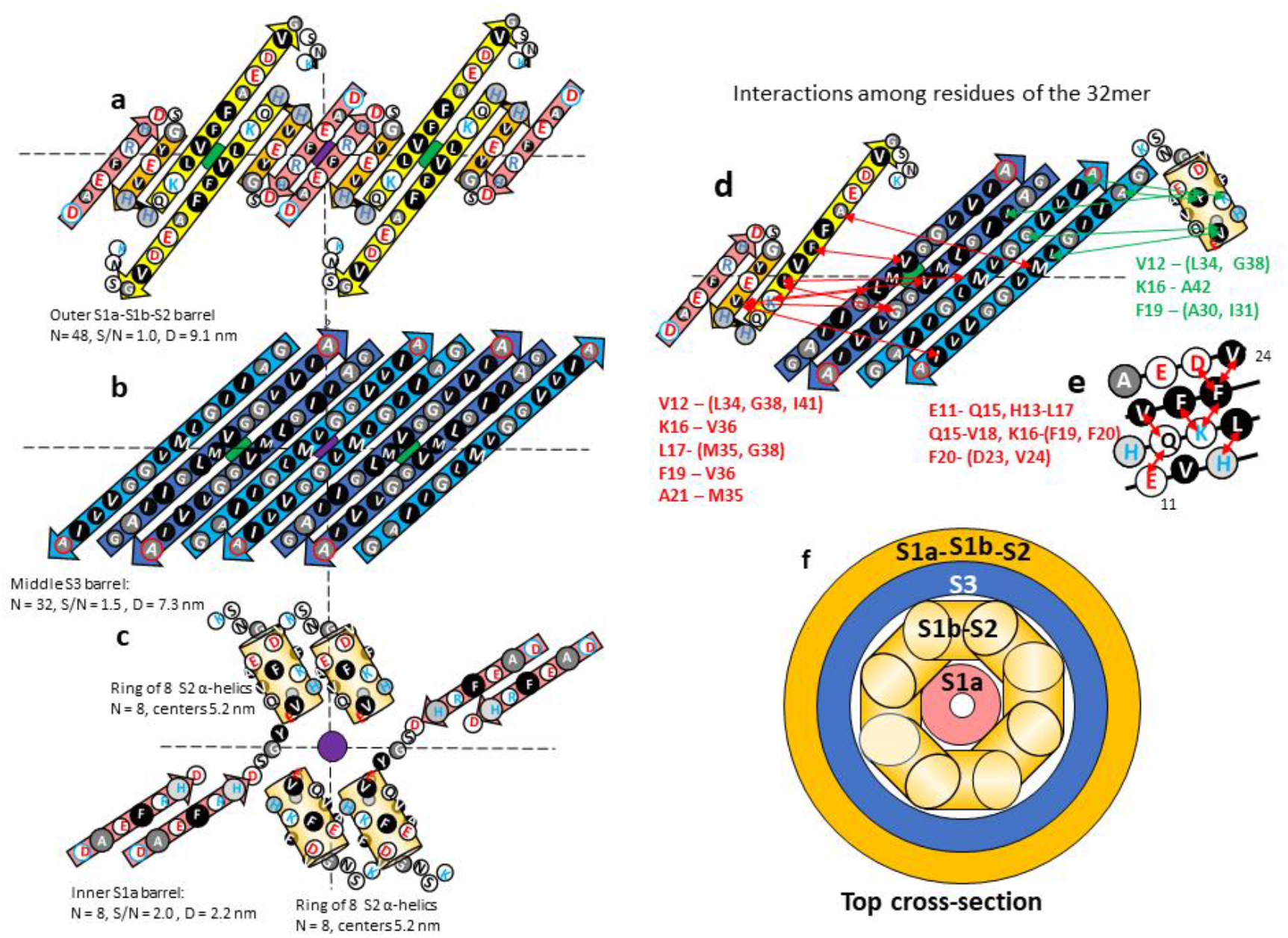
Model of the 32mer WCsAPF. (a-c) Flattened representation of β-strands for four subunits. The green and purple circles behind the strands indicate the axes of 2-fold perpendicular symmetry and the horizontal dashed line indicates the plane containing those axes. Parameters of the β-barrels are indicated beside or below the schematics. (a) The 3-stranded S1a-S1b-S2 β-sheets that comprise the outer β-barrel. (b) The S3 β-strands of the middle β-barrel. C_out_ S3 strands are darker blue. (c) The S1b-S2 α-helices and S1b β-strands of two TIM barrel-like structures of C_in_ subunits. (d) Interactions indicated by NMR studies of Gao et al.^32^ between residues of S1b-S2 and S3. Red lines and residue list on the left side indicate possible interactions between S1b-S2 residues and outwardly-oriented S3 side chains; green lines and residue list on the right indicate possible interactions between the C_in_ S1b-S2 α-helix residues and inwardly-oriented S3 side chains. (e) A helical net representation of the S1b-S2 α-helix showing observed interactions among residues that are sequentially separated by three or four positions. (f) Schematic cross-section of the upper portion of the 32-mer model.

Prediction of the structure of the S1-S2 portion of C_in_ is more difficult. Gao et al.^32^ developed a method to stabilize and isolate 150-kDa Aβ42 oligomers, which they conclude are composed of 32 subunits. These oligomers may correspond to the 32mer WCsAPF of Fig. 7. Their solid-state NMR studies indicate that S3 forms antiparallel β strands centered at V36, consistent with our model and the structure of the central S3 strands of the tetramer. Their results for S2 strands are more complicated. Based on their analysis of chemical-shifts for backbone C sites, they concluded that residues 11-24 have a β secondary structure, but that these residues occupy multiple magnetically inequivalent sites. Additional NMR results indicated that proximal residues are sequentially either three or four positions apart on S1b-S2. They interpreted these data as indicative of out-of-register parallel β-sheets in which the strands are shifted three positions relative to one another. This arrangement is not possible if S2 segments form antiparallel β-barrels.

An alternative possibility is that the assembly contains two distinct S2 conformations, one that has β-secondary structure and another that has α-helical secondary structure in which residues three or four positions apart are proximal (Fig. 8e). If so, then some results may come from the β-structure while other results come from the α-conformation. We propose that S1b-S2 α-helices of C_in_ monomers comprise the inner ring of the 32mer sAPFs. These helices are substantially shorter than the β-strands and stack end-on inside the S3 barrel. Although Gao et al.^32^ concluded that S1a segments are disordered, they may form an inner, parallel β-barrel for C_in_, and part of an antiparallel β-barrel for C_out_. If so, the portion of the assembly inside the S3 β-barrel would resemble a TIM αβ-barrel in which eight parallel α-helices surround an eight stranded parallel β-barrel. This motif has been observed in numerous soluble proteins^60^ and could contribute to the stability of 150-kDa oligomers. A high stability of this structure could also explain why the putative 32mers are the most frequently observed WCsAPFs (Fig. 7).

Additional NMR results of Gao et al.^32^ regarding interactions between S1b-S2 residues and S3 residues are consistent with our model if one assumes some data are due to interactions with C_out_ and others with interactions with C_in_ (Fig. 8). Note that the three residues (V12, K16, and F19) proposed to interact with S3 residues near the beginning and ends of S3 strands are on the same face of the putative α-helix. In spite of the relatively consistent agreement between this model and the NMR data, we cannot be confident any of the putative 32mer APFs proposed here correspond to the 150-kDa oligomer studied by Gao et al.^32^ because different procedures and conditions were used to obtain the structures.

An atomic-scale model of a 32mer sAPF is shown in Fig. 9. Side chains of the S1b-S2 helix make direct contact with side chains of S3 β-strands indicated in green in Fig. 8. Some side chains of the S1b and S2 β-strands predicted by NRM to interact did not make direct contact, but distances between the closest side chain atoms were relatively small. One interaction proposed by the NMR study, between K15 and M35, was not satisfied. The EM images of the putative 32mer in Fig. 7 closely corresponds to the size and shape of the cross-section pictures of Fig. 9e and f (also see Supplement Fig. S4). The atomic-scale model is also consistent with our modeling criteria: almost all conserved hydrophobic side chains are buried and pack tightly in at least one of the two conformations, almost all charged groups form salt-bridges, and almost all backbone polar atoms form H-bonds due to the high content of β and α-secondary structure.

**Figure 9.**
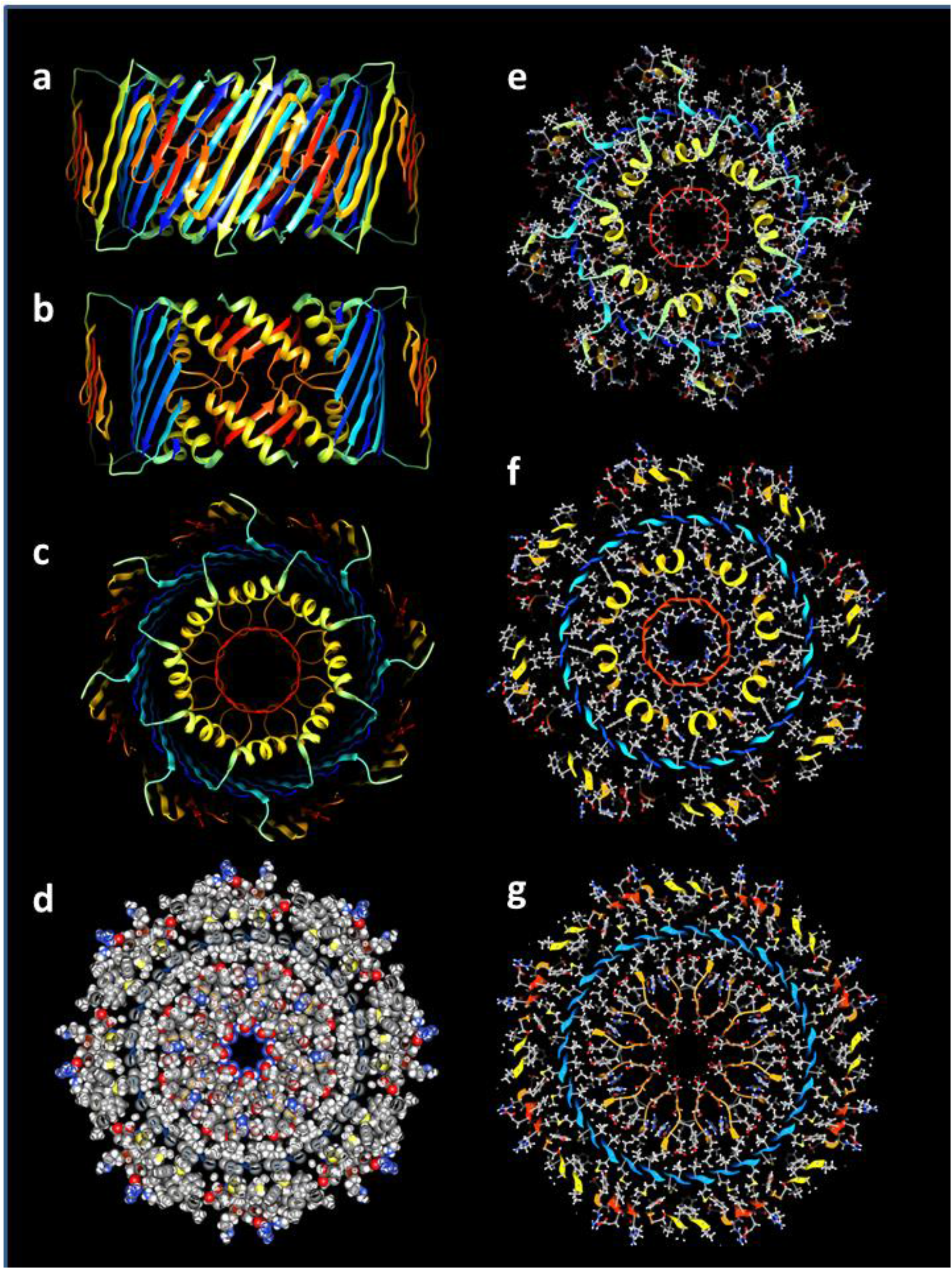
Atomic-scale model of the 32mer WCsAPF. (a-c) Rainbow-colored ribbon representation of the backbone. (a) View from the side. (b) Same as (a) except the outer and S3 strands have been clipped away to show the inner S1b-S2 helices surrounding the S1a 8-stranded β-barrel. (c) View down the central axis of the assembly. (d) Sphere representation of side chains colored by element for a central cross-section illustrating tight side chain packing. (e-g) Stick and ball side chains colored by element and backbone ribbon for the (e) top, (f) slightly lower, and (g) middle cross-sections.

#### DCsAPFs and their relationship to bAPFs

The second category of sAPFs, DCsAPFs, have a dark outer ring with cogs surrounding a light ring, which surrounds a dark circular center. Unlike the white-centered sAPFs, DCsAPFs with the same number of cogs can have different diameters (Fig. 10). Also, WCsAPFs and DCsAPFs of the same approximate diameters often have differing number of cogs and thus different symmetries; e.g., the putative 32mer WCsAPF has eight cogs and 8-fold radial symmetry, whereas the putative 32mer DCsAPF has four cogs and 4-fold radial symmetry. Comparison of sizes and shapes of DCsAPFs to those of bAPFs suggest that the number and distribution of DCsAPF cogs are like those of the beads of bAPFs. Most DCsAPF appear to have morphed from bAPFs composed of the two smallest sizes of beads; i.e., the ones we propose to be hexamers and octamers (outlined in purple and red in Figs. 2 and 10). If so, then the radial unit-cells should have six and eight monomers for the two principal groups of DCsAPFs.

**Figure 10.**
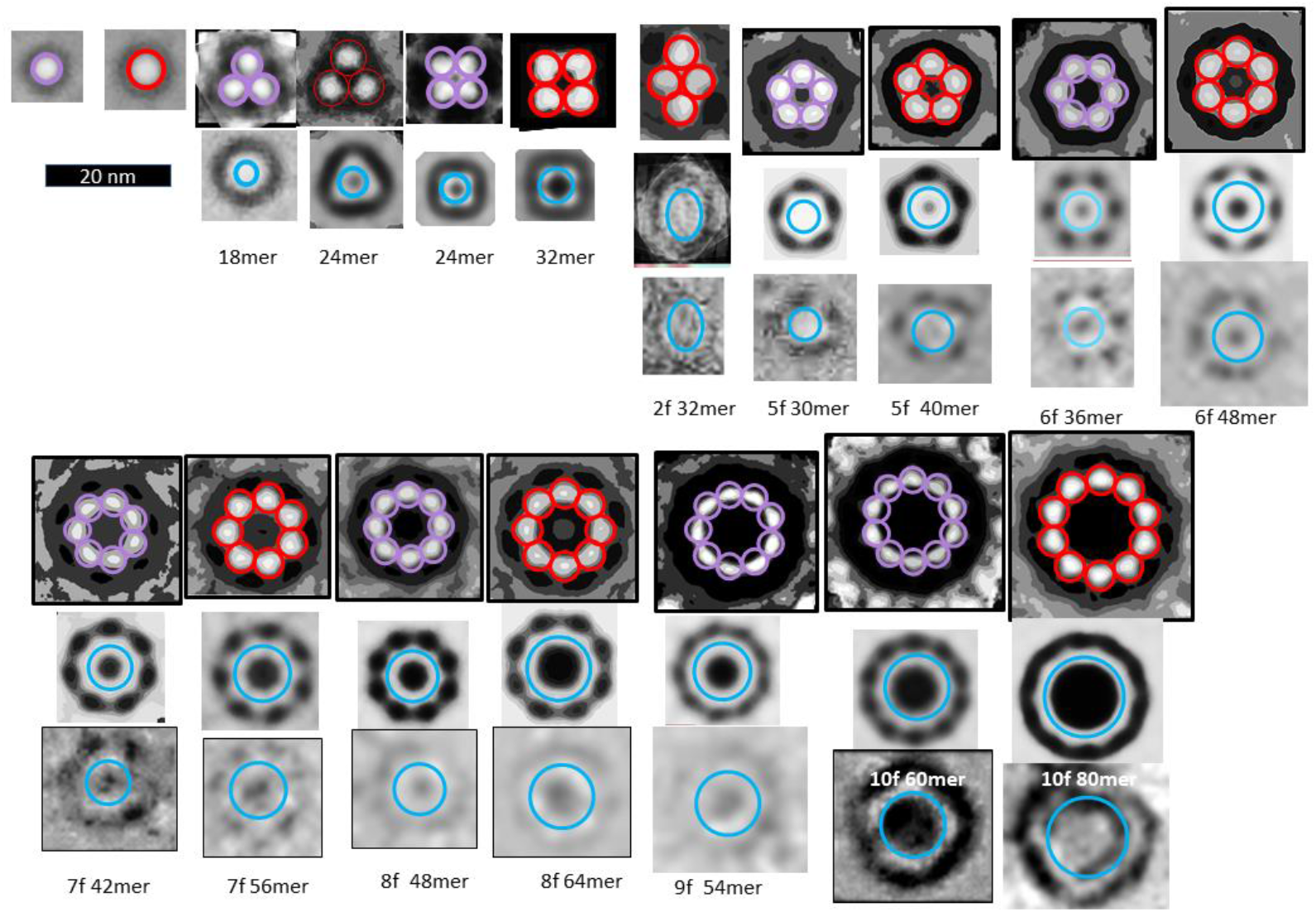
Comparison of DCsAPFs images to bAPF images of Fig. 2. The bAPF images are those proposed to be formed by hexamers (purple circles) or octamers (red circles). Averaged and individual BCsAPF images are illustrated in the two rows below the bAPF images. The blue circles near the middle of the light-shaded rings of the BCSAPFs have diameters predicted for walls of S3 β-barrels (if N is the number of monomers proposed for the APF and S/N = 1.0 for 18-32mers, 2/3 for APFs developed from hexamers, and 3/4 for APFs developed from octamers).

The blue circles superimposed on the white rings of the DCsAPFs in Fig. 10 have the diameter predicted for a S3 β-barrel with the number of strands suggested by the number of monomers proposed for the associated bAPF; e.g., the bAPF composed of six small beads (hexamers) should have 36 monomers and the S3 β-barrel of the associated DCsAPF should have 36 β-strands. S/N values for these putative S3 β-barrels are 1.0 for the 18-32mers, and 2/3 or 3/4 for larger multiples of hexamers or octamers. Thus, these DCsAPFs are consistent with the hypothesis that an antiparallel S3 β-barrel is surrounded by a S1a-S1b-S2 β-barrel in which the S/N values change in a manner that maintains a gap distance between the walls of the barrels near 1.0 nm. It is difficult to predict based on these images whether there is a S1-S2 β-barrel inside the S3 barrel because the interior is completely dark. If the dark center is due to the presence of hexane, then hexane may shield the inner surface of the S3 β-barrel from water and any interior S1 and S2 segments may interact with the surface of a hexane layer.

An interesting exception to the general rule of relatively circular sAPFs are the putative oval-shaped 32mers proposed to have developed from diamond-shaped clusters of four octamers (7^th^ image of Fig. 10). Seven such images were identified in the portion of the micrograph of Fig. 1b that contains the WCsAPFs. These were the only oval sAPFs observed in this region of the EM and they all have similar sizes and shapes, and, like WCsAPFs, they have alternating dark and light layers with a white center. In this instance the sAPFs may have retained the general shape of its ancesteral bAPF rather than adopting β-barrel structures with circular cross-sections. The other possible exception are the DCsAPFs we propose to have 10-fold symmetry. Some of the isolated images appear to have 5-fold symmetry, which when radially averaged with 10-fold symmetry, would appear to have ten cogs. 5-fold symmetry could result if each radial unit-cell has 12 or 16 monomers instead of six or eight, which could occur if they developed from bAPFs composed of five dodecamers or hexadecamers.

#### SsAPFs

Many sAPF images have no discernable cogs. These super smooth APFs (SsAPFs) simply have light-shaded rings with darker shading on their exterior and interiors. However, some do appear to have radial symmetry, as indicated by the radially-averaged images. Our working hypothesis is that cogs appear when triple-stranded S1a-S1b-S2 comprise the outer β-barrel because the S1a and S1b strands are much shorter than the S2 strands (Fig. 6), and that cogs do not appear when S1-S2 hairpins comprise the outer barrel because the continuous S1 strand is about the same length as the S2 strand (Fig. 11). There are two mechanisms by which the outer and inner S1-S2 β-barrels can have larger and smaller diameters than those of the central S3 β-barrels. In most cases the numbers of strands in the outer and inner barrels are likely greater and less than those of the S3 barrels. This situation could develop from bAPF precursors if S1-S2 strands inside the bAPF ring become inner barrel strands and those outside the ring become outer barrel strands. As the SsAPF’s increase in size the ratio of outer to inner barrel strands should decrease, and for very-large SsAPF’s it likely becomes 1.0. In these large SsAPFs the S/N values of the outer and inner barrels likely become greater or less than those of the S3 barrels. Sizes of the SsAPF vary greatly, and many larger ones are less circular than those illustrated in Fig. 11.

**Figure 11.**
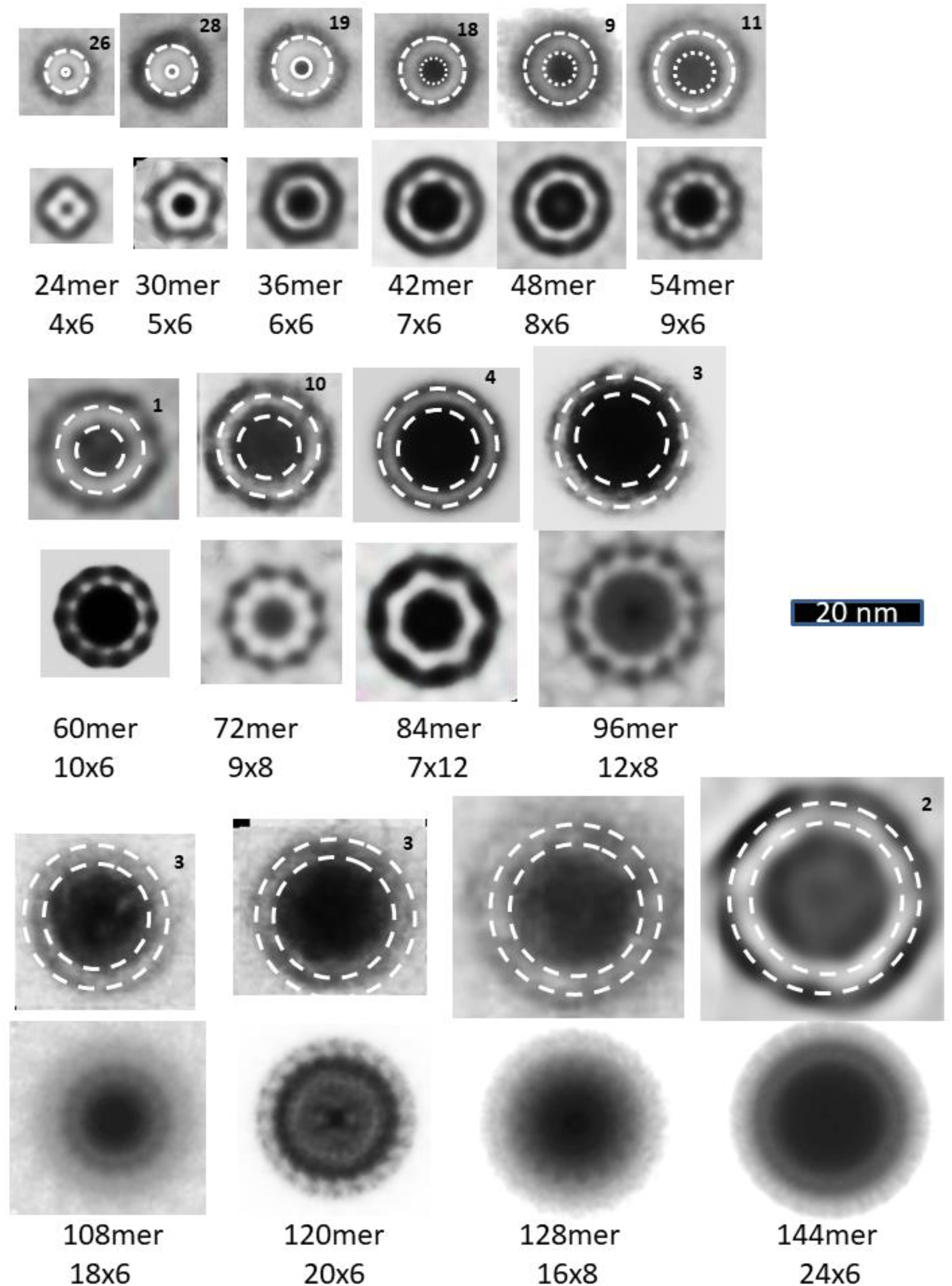
Images of SsAPFs averaged without and with radial symmetry. The top rows of circular SsAPFs images were generated by averaging multiple images of the same size but without radial averaging; the number of images that were averaged is indicated in the upper right corner. Dashed circles represent approximated edges of the SsAPFs walls; the gap distance between the two circles is 3.0 nm. The images below these were obtained from individual images that were radially averaged. The proposed numbers of monomers, of radial unit-cells, and of monomers/unit-cell are indicated below the images.

The thickness of the light rings of SsAPF’s appears to be between 3 and 4 nm regardless of the size or shape of the SsAPF. This thickness is consistent with three or four β-sheet layers that are each about 1 nm apart.

## SUMMARY AND CONCLUSIONS

This is the second paper of a series on the concentric β-barrel hypothesis for non-fibril amyloid assemblies; the first was on Synucleins, especially α-synuclein associated with Parkinson’s Disease^42^. These papers are follow-ups to and expansion of our earlier publications on concentric β-barrel models of Aβ42 oligomers, APFs, and ion channels^22,23^, utilizing recent experimental results and image averaging. Some may consider Aβ42 APFs of secondary interest because they are not toxic and do not form transmembrane channels^39^. Nonetheless, we have chosen to emphasize them because they provide the best pictorial evidence we have for concentric structural architectures. They also provide clues to the structure of smaller oligomers, which are thought to be important in Alzheimer’s disease and may be precursors to transmembrane channels.

Hypothetical models and concepts are vital for research because they may suggest experiments that otherwise would not be performed or funded. Alternative models such as those described in the Supplement are useful because they make different predictions that can be tested. For example, the same residue of adjacent monomers interacts at axes of 2-fold perpendicular symmetry, but the predicted interactions differ among models. Introduction of a cysteine residue at these positions could create disulfide-linked dimers that stabilize some assemblies while preventing formation of others. Doing so could reduce the polymorphism, reduce disorder that complicates structural studies of amyloids, and help identify which models are better. Likewise, models of oligomers can be used to identify possible target segments to which binding of antibodies or drugs could preclude farther growth of the assembly. Thus, having a basic understanding of how amyloids develop and morph among many experimentally observed configurations may provide new insights for development of molecular preventions, treatments, and possibly cures of the devastating diseases associated with amyloid misfolding.

## Authors’ contributions

Most of the theory and graphics were developed by HRG, who wrote most of the text. SRD developed the β-barrel software and co-created the 3D versions of the models, and provided valuable suggestions for the theory, figures, and editing of the manuscript. RK performed the EM studies of the annular protofibrils and provided the EM images.

## The authors have no competing interests

Original EM images will be supplied by HRG at upon request.

