## Supplement figures for "The amyloid concentric β-barrel hypothesis: Models of amyloid beta 42 oligomers and annular protofibrils"

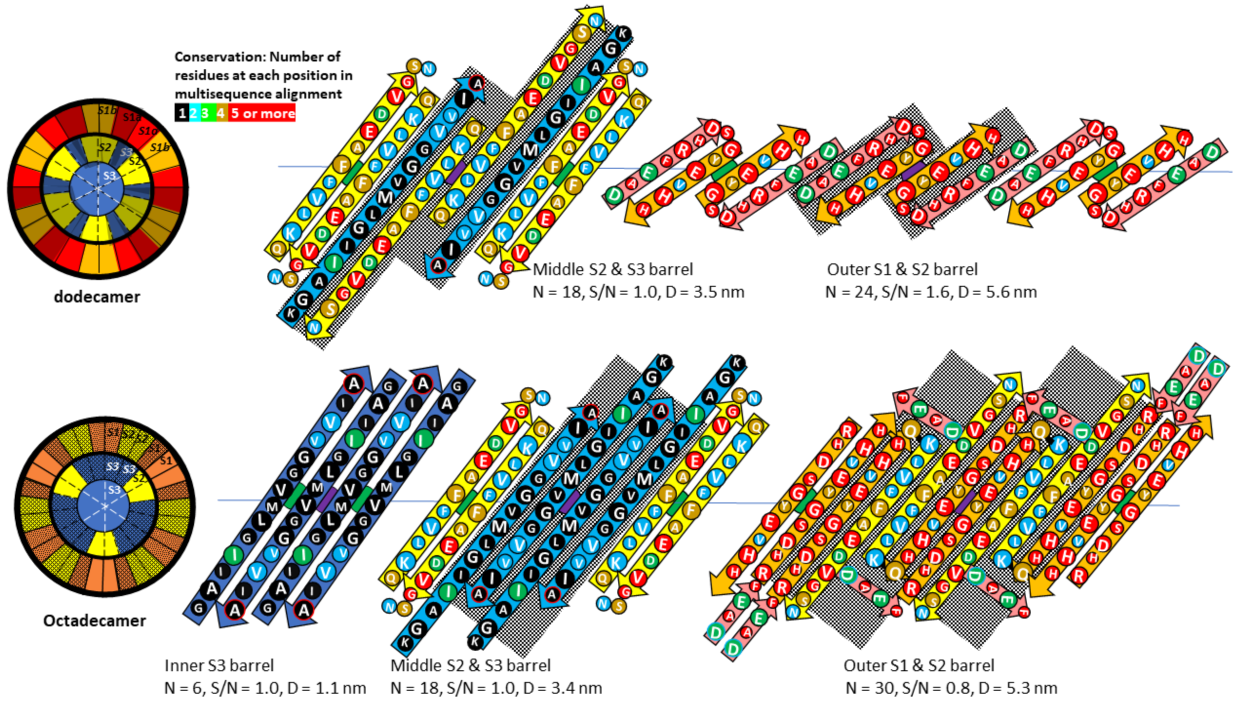


Figure S1. Alternative models for the dodecamer and octadecamer. Additional subunits that are added to the six-subunit core are shaded. (Dodecamer) The relative positions of the S2 strands and S1a-S1b hairpins differ from the model in Fig. 4 of the text. (Octadecamer) The middle barrel is similar to that of the dodecamer except that the central S2 strands of the additional subunits are replaced by S3 strands. All strands intermesh at the perpendicular axes of 2-fold symmetry. All S1 strands are continuous and are components of the outer barrel, which has 30 strands instead of 24 of the text model. H6 is adjacent to H15 where they may comprise a $Cu^{2+}$ binding site [38] and two Y10 residues are adjacent at a perpendicular axis of 2-fold symmetry where they may cross-link under oxidizing conditions to form dimers [39]. The first three or four residues of S1a are not part of the S1-S2 β-hairpins but their relatively conserved D1 (N-terminus) and E3 residues (green) may interact favorably with relatively conserved Q15, K16, D23, K28, A42 (C-terminus) and with each other.

```
DAEFRHDSGYEVHHQKLVFFAEDVGSNKGAIIGLMVGGVVIA  42
ETKYQREAAFDLRPERQMLLPDEMSTK   V       II
NSDEHQSTP-Q YDK    YTKNSNP    L
 P VGLNVSIA S-R    ISN LPF
 I DKD--E G NR       Q AR
 K GEMG N
   MTVT -
   A -  Q
   N    R
        D
```

**Number of residue types at each position:** 1 2 3 4 5 or more

Figure S2. Results of multisequence alignment of 2500 Aβ homologs that includes most vertebrates from bony fish through mammals. Sequences that appeared to be partial were excluded, but over 2000 remained. The top line is the sequence of human Aβ42 colored by the number of residue types at each position of the alignment. Residue types and deletions that occur in other sequences are indicated below the human sequence.

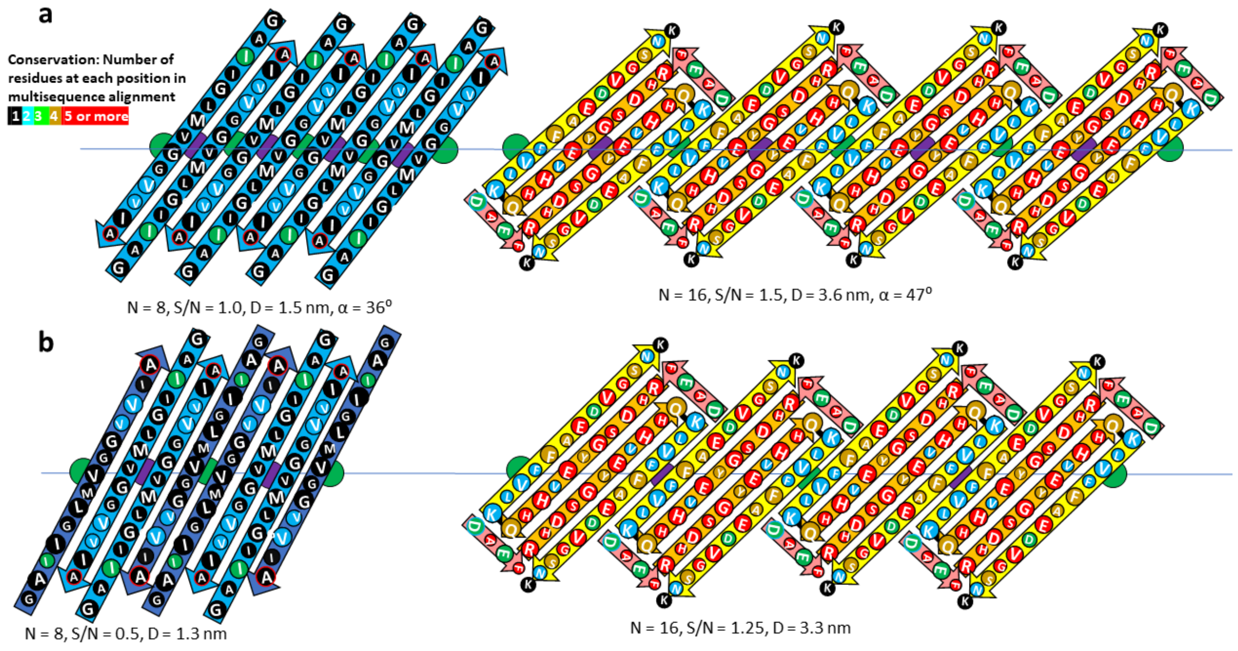


Figure S3. Flattened representation of alternative octamer models of Aβ42. The core 8-stranded S3 β-barrel has two conformations like those of the tetramer [32] and proposed for the WCsAPFs. Those with odd numbered side chains oriented outwardly have lighter blue arrows. The antiparallel pairs of S1-S2 strands of the outer barrels have conformations like those proposed in Fig. S1 for the octadecamer. The two models differ from each other primarily in the tilts (S/N values) of the strands and diameters of the barrels. The models differ from that of text Fig. 6 in the orientation of half the S3 segments and in the relative positions of S1 and S2 strands.

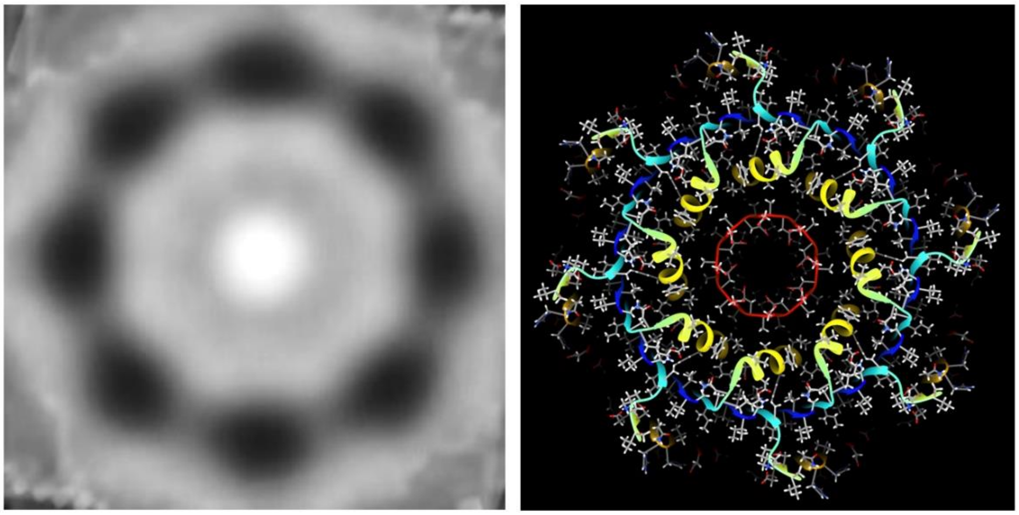

Figure S4. Comparison of averaged EM image of the putative 32mer WCsAPF to a cross-section of the atomic scale model. Both are to the same scale.

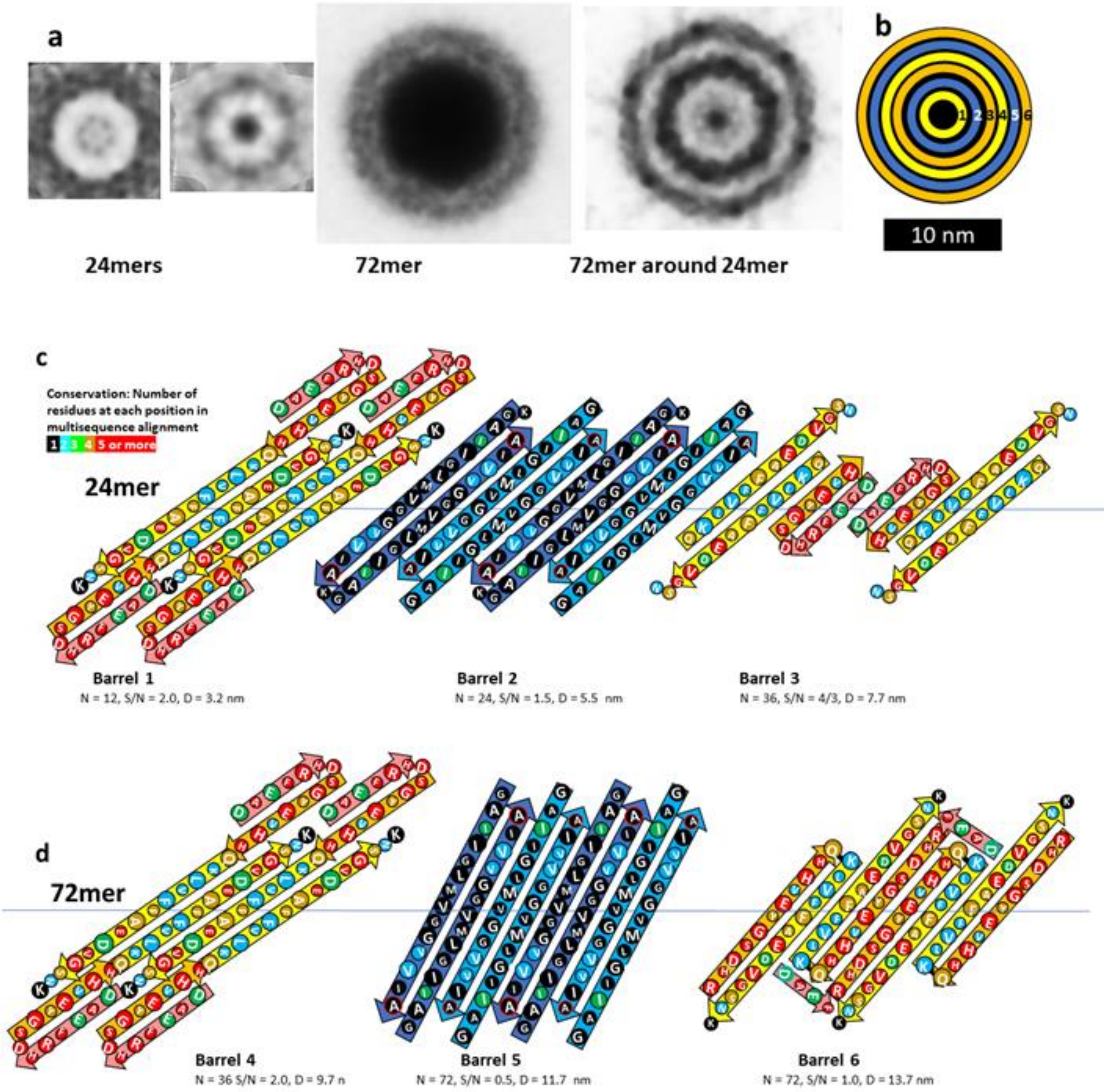


Figure S5. Averaged EM images and models of the 72 around 24mer sAPF. (a) Comparison of sAPF EM images of a WCsAPF 24mer, a DCsAPF 24mer, a DCsAPF 72mer, and the putative 72mer-around-24mer sAPF. All were averaged with 6-fold radial symmetry. (b) Schematic cross-section representation of the model's six concentric β-barrels. The concentric rings have diameters predicted by the models in c and d. (c & d) Flattened representation of eight monomers for the 24mer and 72mer. Side chains are colored by conservation among Aβ homologs. (a: 24mer) The inner 12-stranded barrel is formed by $C_{in}$ S2 strands with possible S1a-S1b β-hairpins in series. Half of the 24 S3 strands of Barrel 2 have the $C_{in}$ conformation and the other (darker) half have $C_{out}$. S1a-S1b-S2 β-sheets of $C_{out}$ subunits comprise the 36-stranded Barrel 3. (d: 72mer) S2 strands and S1a-S1b β-hairpins of Barrel 4 have the same structure as in Barrel 1 except that the barrel has three times as many strands. The 72-stranded S3 Barrel 5 resembles Barrel 2 except that the strands are less tilted (S/N = 0.5 instead of 1.0). Adjacent pairs of S1-S2 β-hairpins of Barrel 6 have structures resembling those proposed in Fig. S1 for the dodecamer and Fig. S3 for the octamer. The predicted gap distances are ~ 1.1 nm between Barrel 1 and 2 and between Barrel 3 and 4 and are ~1.0 nm for the remaining barrels.
